# Multiplexed amplicon sequencing reveals the heterogeneous spatial distribution of pyrethroid resistance mutations in *Aedes albopictus* mosquito populations in Southern France

**DOI:** 10.1101/2023.08.05.552101

**Authors:** Albin Fontaine, Antoine Mignotte, Guillaume Lacour, Agnès Nguyen, Nicolas Gomez, Lionel Chanaud, Grégory L’Ambert, Sébastien Briolant

## Abstract

The risk of mosquito-borne diseases transmission is moving fast toward temperate climates with the colonization and proliferation of the Asian tiger mosquito vector *Aedes albopictus* and the rapid and mass transport of passengers returning from tropical regions where the viruses are endemic. The prevention of major *Aedes*-borne viruses heavily relies on the use of insecticides for vector control, mainly pyrethroids In Europe. High-throughput molecular assays can provide a cost-effective surrogate to phenotypic insecticide resistance assays when mutations have been previously linked to a resistance phenotype. Here, we screened for the spatial distribution of *kdr* mutations at a large scale using a two-step approach based on multiplexed amplicon sequencing and an unprecedented collection of field-derived mosquitoes in South of France. We identified the presence of the V1016G allele in 14 sites. The V1016G allele was predominantly found in South-East France close to the Italian border with two additional isolated sites close to Bordeaux and Marmande. All mosquitoes were heterozygous for this mutation and should not be phenotypically resistant to pyrethroid insecticide. Four other mutations were identified in our targeted genomic sequence: I1532T, M1006L, M1586L, M995L. Sequencing a section of maternally inherited mitochondrial genome confirmed that the spread of *Ae. albopictus* in France originated from founders with haplogroup A1. These findings contribute to the broader understanding of resistance dynamics in Europe and can inform targeted approaches to mitigate the impact of resistance on vector control.

---

Once confined to tropical areas, the risk of mosquito-borne diseases transmission is now moving fast toward temperate climates, fostered by the colonization and proliferation of the Asian tiger mosquito vector *Aedes (Stegomyia) albopictus* and the rapid and mass transport of passengers returning from tropical regions where the viruses are endemic. The unusually high secondary autochthonous cases of dengue virus (DENV) infections in South of France in 2022 illustrates the risk and is sounding the alarm^1^. The prevention of major *Aedes*-borne viruses heavily relies on the use of insecticides for vector control. In Europe, deltamethrin (a pyrethroid insecticide) is the only insecticide authorized in space spraying to target flying adult mosquitoes^1–3^. Resistance toward this insecticide has been described in *Ae. albopictus* populations throughout the world, including Europe^3–5^, but limited information is yet available for France. Their spread can negatively impact the effectiveness of vector control interventions and put in jeopardy our very limited defense line.

Monitoring phenotypic insecticide resistance at a large scale is expensive, time-consuming, and laborious. High-throughput molecular assays can provide a cost-effective surrogate when mutations have been previously linked to a resistance phenotype. In addition, molecular methods can detect resistance alleles before they reach fixation and can thus be used as an early-warning approach^6^. Mutations at 2 codon positions (V1016 and F1534) in the voltage sensitive sodium channel (Vssc) gene were experimentally identified as the main knockdown resistance (*kdr*, the main resistance mechanism to pyrethroids) mechanism in *Ae. albopictus*^4,7^. Here, we report a two-step approach based on multiplexed amplicon sequencing to screen for the spatial distribution of *kdr* mutations at a large scale using an unprecedented collection of field-derived mosquitoes sampled from 95 sites across 61 municipalities alongside a West to East transect in South of France.

## Results

### Screening of KDR mutations in pool DNA amplicons sequencing

A total of 547 mosquitoes collected from a West to East transect in South of France from June 2021 to September 2021 at 95 sites in 61 municipalities, either at the egg or adult stage, were grouped by sites into 100 pools. Two non-overlapping genomic DNA fragments covering 4 exons in the *Vssc* gene (exon19-like, exon20-like, exon27-like and exon28-like, as referred to the JAFDOQ010000349.1 annotation file) were amplified using eight different 6 bp barcodes incorporated at the 5’ end of the forward primers (Supplementary table 2). The combination of barcodes and dual indexing allowed the deep sequencing of 13 *super-pools* instead of the original 100. The sequencing generated an average depth of 12,779 X for amplicon 1 (327 bp, exon19-like and exon20-like) and 3,336 X for amplicon 2 (500 bp, exon27-like and exon28-like), per pool after demultiplexing.

A total of 651 mutations were detected on the target region of the *Vssc* gene with allele frequencies ranging from 0.1% to 99.9% (median: 3.7%, 1^st^ quartile: 0.3%, 3^rd^ quartile: 1.4%) (Supplementary figure 1) across pools. A total of 445 mutations were located on exons, among which 131 (29%) were synonymous and 314 (71%) non-synonymous. These non-synonymous mutations were located at 304 unique positions and had an overall low allele frequency with a median of 0.33% (1rst quartile: 0.25%, 3^rd^ quartile: 0.51%) across pools (Supplementary table 3). Seventeen of them (5.4%) had mean allele frequencies > 2% across pools (Supplementary figure 2). Mutations M1006L and I1532T, detected in 98 and 74 pools, respectively, were one of the most prevalent.

KDR V1016G and V1016I mutations were detected in 19 and 3 pools, respectively (Figure 1-A). Pools with mutation V1016I had very low allele frequencies (below 0.625%, which is the theoretical frequency threshold if one heterozygote allele is detected in the biggest pools of N=80). KDR V1016G mutation was preferentially detected in the Southeast of France from Marseille to Nice with two exceptions in Bordeaux and Marmande (Figure 1-A).

**Figure 1:**
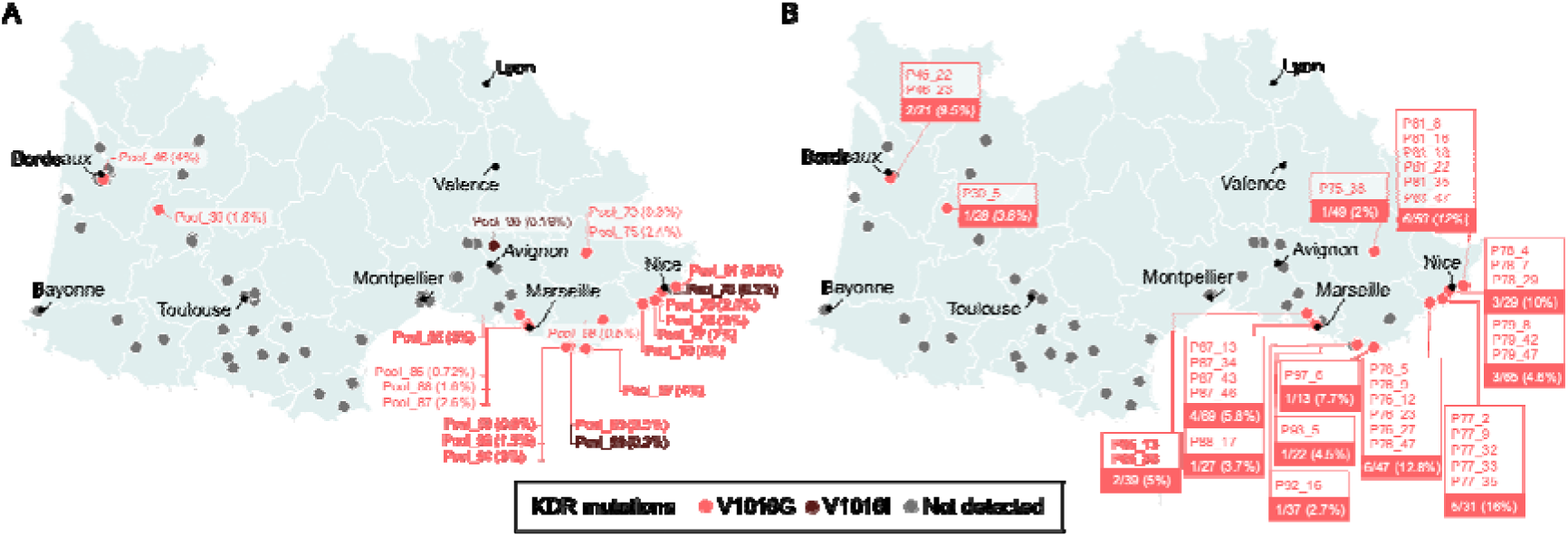
Geographic location of alleles confirmed in knockdown resistance in *Aedes albopictus* in South of France. A) Location and frequencies of KDR alleles as revealed by amplicon sequencing based on sequencing of DNA from pooled mosquito heads. Allele frequencies are represented into brackets for each locality. B) Location and prevalence of KDR alleles as revealed by amplicon sequencing on single mosquitoes from each locality. The identity and prevalence of mosquitoes carrying the mutations are represented for each locality. Mosquitoes are identified based on their original pool number and a unique number. Grey points represent localities where no confirmed KDR alleles were identified.

### Confirmation of KDR mutations in single mosquito DNA amplicons sequencing

Single mosquito DNA sequencing was implemented to confirm mutations revealed by pool DNA sequencing and to determine their prevalences and genotypes (heterozygote / homozygote). Genetic variations were detected at 135 positions over the target regions of the *Vssc* gene. A total of 32 mutations were located on exon-like regions, among which 27 (84%) were synonymous and 5 (16%) non-synonymous: M1006L, M995L, V1016G, I1532T, and M1586L (Supplementary table 3). Importantly, all these mutations were previously identified in the top 20 most frequent mutations in pool DNA sequencing (Supplementary figure 2). However, some mutations identified in pool DNA sequencing were not confirmed when sequencing individual mosquito DNA.

The M1006L mutations was the most prevalent (detected in 215 mosquitoes from 45 sites), followed by the I1532T mutation that was detected in 54 mosquitoes from 16 sites. KDR V1016G mutation was detected in 37 mosquitoes from 14 different sites. This mutation was detected in the same sites than pool DNA sequencing except for pool_73, pool_86, pool_98, pool_90, and pool_94. Allele frequencies for these pools were mainly < 1% and might be attributed to DNA contamination during the DNA extraction procedure. KDR V1016G mutation in single mosquito DNA was not detected in sites where the mutation was not reported by pool DNA sequencing. All mosquitoes were heterozygotes for this mutation. The prevalence of mosquitoes carrying the V1016G mutation ranged from 2% to 16% across sites (Figure 1-B). Single DNA sequencing confirmed the presence of KDR V1016G mutation in Southeast France, close to the Italian border where it has already been described since 2019^3,4^ and in a cluster located in the West in Bordeaux and Marmande.

### Geographical dispersion of mosquitoes carrying KDR mutations as revealed by their mitochondrial DNA

The amplicon-based library targeting the *Vssc* gene was complemented with ligase-based tiling amplicons that amplified a region of the mitochondrial genome in each *Ae. albopictus* mosquitoes. Genomic regions aligning to the targeted mitochondrial genome had a low depth (mean ± SD: 5.8X ± 33), as compared to the *Vssc* gene. A total number of 126 samples out of 1,167 (11%) could be however selected to be included in the phylogenetic analysis with background reference sequences that represented the worldwide diversity of *Ae. albopictus* mitogenomes haplogroups^8,9^. The phylogenetic analysis revealed that all collected *Ae. albopictus* mosquitoes originated from founders with haplogroup A1 (Figure 2). The KDR V1016G allele was found in mosquitoes from different maternal lines. One mosquito from the West (P30_5) carrying the KDR V1016G mutation had a mitochondrial DNA genetically close to mosquitoes from the East (Figure 2), suggesting long-range dissemination of the resistance allele through transports.

**Figure 2:**
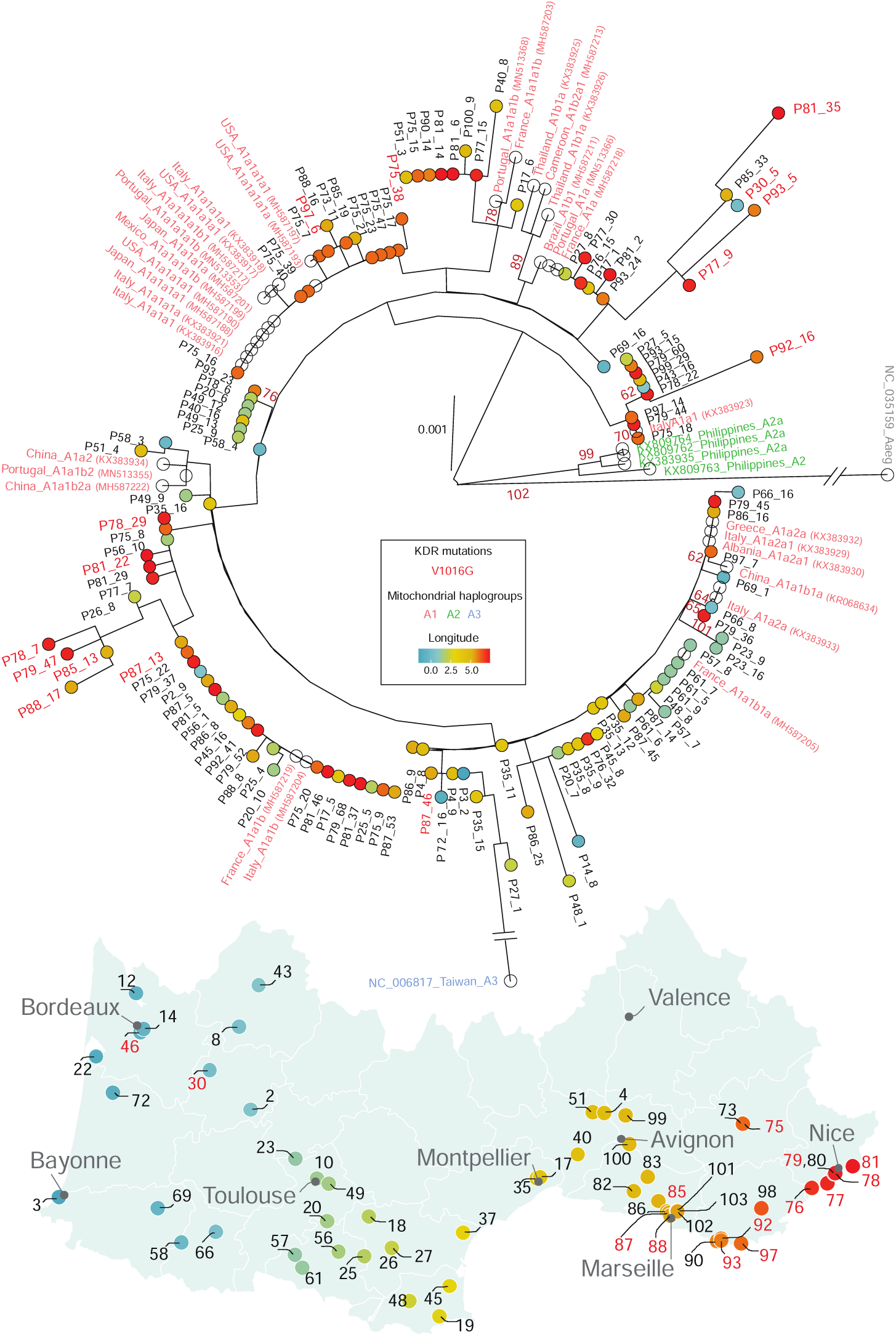
Phylogenetic relationships among a subset of *Aedes albopictus* mosquitoes analyzed in this study based on a curated alignment of 3,243 bp nucleotides region of the mitochondrial genome. The mitochondrial genome of *Ae. aegypti* (NC_035159) was used as an outgroup in the phylogenetic tree. The best-scoring maximum-likelihood (ML) tree was generated with 120 bootstrap replicates. Only bootstrap scores > 60 are represented in dark red on the figure. Mosquitoes are identified based on their original pool number and a unique identifier. Pool localities are represented on a map with a color code representing the longitude (West to East transect gradient is represented with a blue to red color gradient). Localities with at least one mosquito carrying a KDR allele are represented in red on both the map and the phylogenetic tree.

## Discussion

Pyrethroid insecticides are nowadays widely used in agriculture or as indoor/outdoor residual or space spraying for adult mosquito control throughout the world because of their low acute toxicity on mammals and high and fast activity in insects. Mutations in the *Vssc* gene were experimentally identified as one of the major knockdown resistances (*ie*. *kdr*) mechanisms in insects, together with metabolic resistance mainly mediated by P450 monooxygenases^10,11^. KDR mutations were originally discovered on the model organism *Musca domestica*^12^, and mutations found in other insects were named based on the codon position of this house fly reference genome. Several mutations were documented on the Vssc gene in *Ae. aegypti* (*ie.* V410L, S989P, I1011M/V, V1016G/I, I1532T, F1534S/L/C, and D1763Y), but a few of them have been confirmed to be functionally associated with pyrethroids resistant phenotypes (*i.e.*, V410L, S989P, I1011M, V1016G and F1534C)^13^. Some mutation combinations can engender extreme resistance in *Ae. aegypti*, such as the triple mutant 989P/1016G/1534C haplotype^14^. The KDR F1534C mutation was the first to be reported in *Ae. albopictus* in Singapore^7^ in 2011, followed by mutations V1016G/I, and F1534S/L in different parts of the world alone or in combination. The KDR V1016G allele was recently found in *Aedes albopictus* populations from Italie, Vietnam^4^ and China^15^. At the homozygous state, this mutation was shown to confer a higher level of pyrethroid resistance than the previously known alleles, F1534C and F1534S^4^. The KDR V1016G mutation was recently revealed in France in two populations of *Ae. albopictus* from Nice and Perpignan^3^.

Here, the spatial distribution of pyrethroid resistance mutations in *Ae. albopictus* populations in southern France was screened in the most exhaustive sampling work to date in France (95 sampling sites across 61 municipalities), using a two-step multiplexed amplicon sequencing approach. We first implemented a sequencing approach using pooled mosquito DNA per site to reduce the overall sequencing costs. This initial step was able to screen for the presence of KDR mutations in many sites across a wide study area, faster, and with less samples size as compared to a single mosquito DNA screening approach. Several mutations with high allele frequencies and prevalence across sites were detected, including KDR V1016G mutations. Importantly, all mutations subsequently confirmed by single mosquito DNA sequencing were previously identified in pool DNA sequencing. However, some mutations identified in pool DNA sequencing were not confirmed when sequencing individual mosquito DNA (*eg.* V1016I). This can be partially explained by the allele calling program (LoFreq) applied to pool DNA sequencing that is more sensitive to distinguish rare variants than the pipeline applied to single mosquito DNA sequencing. This can create difficulties to distinguish rare variants from sequencing errors^16,17^. The presence of KDR V1016I mutation was not ultimately confirmed by single mosquito DNA sequencing, this can be due to a low allele frequency < 1% in the three pools where it was detected. Pool DNA sequencing allows to identify the sites with the presence of KDR V1016G allele with a perfect sensitivity (100%), albeit not good specificity with 5 sites out of 19 being not confirmed by single DNA sequencing. We suspect that contaminations across samples might had occurred during the grinding step prior the extraction procedure for pool DNA library preparation. This issue can be easily improved in the future. This two steps approach can save time and resources, especially when the presence of the target mutations is anticipated to be scarce, by excluding samples from sites in which the targeted mutations was not detected in a preliminary screening. Efforts and money can then be dedicated in a more efficient way to analyze prevalences and genotypes using single mosquito DNA in selected sites. This method can be readily integrated into routine surveillance programs, allowing for the early detection of resistance before the fixation of mutations and the timely implementation of appropriate control measures.

The V1016G allele was predominantly found in South-East France close to the Italian border with two additional isolated occurrences close to Bordeaux and Marmande. While previous study already reported the presence of the KDR V1016G allele mutation in Nice and Perpignan, our sampling effort across the South of France did not identify any resistance genes in Perpignan. Importantly, genetic resistance to insecticides can be highly clustered even at the small geographic scale. *Vssc* harbouring the V1016G allele was not detected from *Ae. albopictus* collected outside of Hanoi City in Vietnam while it was found in the city^4^. In our study, this mutation was found in population collected in harbor areas in Marseille but not in those collected more inland from the same city. A genome-wide analysis with a high density of nucleic DNA markers revealed a weak genetic structure and high levels of genetic admixture in *Aedes albopictus* populations from Switzerland, supporting a scenario of rapid and human-aided dispersal along transportation routes, with frequent re-introductions into Switzerland from Italian sources^18^.

The use of pyrethroid is strictly regulated in France when there are applied for curative vector control around human cases of dengue, chikungunya or Zika – imported or autochthonous – to reduce the risk of local arbovirus transmission^3^. Paradoxically, there is neither formal prohibition nor any surveillance of the use of pyrethroids by pest control companies for as part of nuisance reduction. The use of insecticides by pest control companies or private individuals might maintain a significant selection pressure on local insect populations. Resistance genes carrying *Ae. albopictus* populations in Nouvelle-Aquitaine sites were not exposed to curative vector control treatment within 150 meters since at least 2020. In contrast, resistance genes were not revealed in mosquitoes from sites which had undergone six repetitions of treatments since 2020. The *de novo* appearance of mutations is a rare event and resistance in a population commonly arises from selection of resistant alleles that are present in a population or from the arrival of individuals with resistance alleles through transport by humans^6,19^. Here, we revealed close genetic relationships between mosquitoes collected in West and East of France that were carrying the V1016G allele using a section of the maternally inherited mitochondrial genome. Altogether, these data suggest that the presence of KDR mutations in France originated from fast transportation between distant populations rather than from *de novo* due to a strong selection pressure.

Although the French Agency for Food, Environmental and Occupational Health Safety (ANSES) established recommendations in 2020 regarding the use of insecticides and the surveillance of resistance in French populations, there is currently no national surveillance program in place^20^. While resistance of vector mosquitoes has been well-documented in overseas territories^11,21,22^, it remains poorly studied in metropolitan France. The presence of insecticide resistance alleles in *Ae. albopictus* populations from different sites in France highlights the need for a continued monitoring of insecticide susceptibility at a wide geographic scale, together with the development of alternative vector control strategies to alleviate the selection pressure. All mosquitoes carrying the V1016G mutation in France displayed a heterozygous genotype. Fixation of KDR V1016G allele, and thereby the occurrence of phenotypic insecticide resistance, can arise rapidly in the presence of a strong selection pressure in areas where the allele is detected even at a low prevalence. There is thus a critical need for the implementation of a comprehensive national surveillance program to monitor resistance spatially and temporally in *Ae. albopictus* populations. Such a program would provide valuable insights into the prevalence and spread of resistance, allowing for timely and targeted interventions to maintain the efficacy of vector control measures. This may include reducing treatments, alternating authorized insecticides over space and time, employing complementary methods such as trapping and innovative control strategies^23,24^ to proactively respond to changes and mitigate the spread of resistance, thereby safeguarding the effectiveness of vector control interventions and protecting public health.

Four other mutations (*ie.* I1532T, M1006L, M1586L, M995L) were identified in our targeted *Vssc* gene sections in this study. Among these 4 mutations, the I1532T was reported in different *Ae. albopictus* populations from Asia^15,25^, Italy^26^ and Greece^27,28^. This mutation was found in mosquito populations from Rome with a high frequency (19.7%) but not in populations collected 570 km away from this city^26^, which further highlight the patchy distribution of *Ae. albopictus* throughout the territory, even at a small geographic scale. Further work is needed to functionally validate or invalidate the impact of M1006L, M1586L and M995L on insecticide resistance.

## Conclusion

Our study provides insights into the spatial distribution of pyrethroid resistance mutations in *Ae. albopictus* populations in the South of France. Here, we demonstrated that pooled-DNA amplicon sequencing can help to reduce the surveillance costs by detecting the presence of known mutations when they are expected to occur at a low prevalence, prior to screen mosquitoes individually. The use of multiplexed amplicon sequencing, with its ability to screen pooled samples and subsequently confirm findings through individual mosquito DNA sequencing, is a valuable tool for monitoring the spatial distribution of resistance mutations. The detection of the KDR V1016G allele in different French localities emphasizes the need for ongoing monitoring and proactive resistance management strategies. These findings contribute to the broader understanding of resistance dynamics and can inform targeted approaches to mitigate the impact of resistance on vector control efforts.

## Materials and methods

### Field-collected mosquitoes

*Aedes albopictus* mosquitoes were collected from the field either at the egg stage using egg-laying traps or at the adult stage using BG sentinel (BGS, Biogents AG) traps at 95 sites in 61 municipalities alongside a West to East transect in South of France from June to September 2021. Adult mosquitoes were captured over one week with carbon dioxide provided as a mosquito attractant and identified morphologically. Mosquito’s eggs from 181 ovitraps were hatched and reared in laboratory until the fourth instar larvae; 2833 larvae were transferred by sites and sampling date into 90% ethanol. All samples were stored at −20°C until the DNA extraction procedure. Traps were mainly placed at hospital, airport, or seaport sites.

### Mosquito DNA extraction

A two-steps approach was implemented to screen for KDR alleles in *Ae. albopictus* mosquitoes: *i*) an initial screening by sequencing pooled mosquito DNA in each site followed by *ii*) sequencing individual mosquito DNA to determine KDR allele prevalence and genotype. A total of 3 to 80 (mean=24.5, SD=15) mosquitoes were selected by site and grouped into 100 different pools. Heads from larvae or adult mosquitoes were dissected under magnifying glasses. Each pool was made up of mosquito heads sampled at the beginning and the end of the sampling period for each site when possible. All mosquitoes from sites in which KDR alleles were detected in step *i* were selected for single mosquito DNA sequencing, excluding damaged mosquitoes. This second selection also included sites without detection of KDR alleles in step *i*, with a total of 56 sites throughout 50 municipalities. Mosquito heads or bodies were grinded in a 96 wells plate using a TissueLyser (Qiagen) for 2 min at 30 oscillation/s. Genomic DNA was then extracted from homogenates using the NucleoSpin 96 Tissue Core Kit (Macherey-Nagel) and stored at −20°C until use.

### Amplicon-based sequencing

We devised an amplicon-based approach that captured 3 main mutations previously reported to be associated with pyrethroid resistance in *Aedes* spp. mosquitoes: S989P, V1016I/G and F1534C/L/S^13,29^. Two non-overlapping amplicons of 327 bp and 500 bp were used to amplify two sections of the voltage sensitive sodium channel (vssc) gene that was mapped on the *Aedes albopictus* isolate FPA chromosome 3 chr3.142 whole genome shotgun sequence (AalbF3 genome assembly, GenBank: JAFDOQ010000349.1). This sequence was identified in the AalbF3 genome assembly based on its genetic homology with *Ae. aegypti* LOC5567355 vssc gene sequence. The first and second amplicon mapped to JAFDOQ010000349.1 reference sequence at positions 1,806,101 to 1,806,578 bp and 1,851,149 to 1,851,765 bp, respectively. Both amplicons covered four exons in the vssc gene: exon19-like, exon20-like, exon27-like and exon28-like. Both targeted genomic regions were amplified in a single reaction to generate sufficient templates for subsequent high-throughput sequencing. Multiplex PCR reactions were performed with 5 μl of purified DNA in a 20 μl reaction mixture made of 5 μl of Hot START 5X Hot Firepol DNA Polymerase mix (Dutscher, France), 1 μl of forward and reverse primers mix at 10 μM (4 μl for 4 primers) (Supplementary table 1), and 11 μl of water. The thermal program was: 10 min of polymerase activation at 96°C followed by 35 cycles of (i) 30 sec denaturing at 96°C, (ii) 30 sec annealing at 62°C and (iii) 1 min extension at 72°C, followed by a final incubation step at 72°C for 7 min to complete synthesis of all PCR products. Illumina Nextera® universal tails sequences were added to the 5’ end of each of these primers to facilitate the library preparation by a two-step PCR approach. Our multiplexing design involves a same barcode inserted in both forward primer’s sequences on each row of a 96 well plates, so that 10 μl of amplified products could be pooled per column (i.e., 8 samples were pooled into a single tube with a final volume of 80 μl). This multiplexing scheme allow a 8-x sample reduction with 96 samples from one plate being grouped into 12 different tubes, or one plate row (Supplementary figure 3).

The individual mosquito KDR library was complemented with a ligase-based tiling amplicon sequencing method to amplify a 4,438 nucleotides region of the mitochondrial genome in each *Ae. albopictus* mosquitoes. The method generates overlapping amplicons of ∼500 base pairs from two multiplexed PCR reactions with 6 primers pairs in each reaction (Supplementary table 1) to generate sufficient templates for subsequent high-throughput sequencing^30,31^. The Hot START 5X Hot Firepol DNA Polymerase (Dutscher, France) add an adenosine nucleotide extension to the 3′ ends of each replicated DNA strands to create an A overhang, which make the product suitable for ligation with T-tailed DNA adaptors. Eight universal barcoded T-tailed DNA adaptors were made by annealing upper and lower oligonucleotides (Supplementary table 1) at 25M in 1X TE and 3M NaCl buffer, starting with 1 min step at 95°C and a constant temperature reduction of −0,1 °C/sec until to reach 12°C. Each T-tailed DNA adaptors integrated one of the 8 barcodes used in the KDR library preparation. One microliter of T-tailed DNA adaptors diluted to 1.5 µM in water was added to 5 µl of amplicons diluted to 1/10 in water and 5 µl of 2X Blunt/TA Ligase Master Mix (New England Biolabs, Herts, UK) and incubated 30 min at 25°C for ligation. No DNA purification was done purposely prior the ligation step to reduce library costs. Ten microliters of adapter ligated amplicons were mixed to 1 uL of KDR library previously diluted 1/10 in water to obtain a KDR/mitochondrion (primer pool 1 and 2) library ratio of 2, based on DNA concentration determined by Qubit fluorometer and Quant-iT dsDNA Assay kit (Life technologies, Paisley, UK) from a random subset of samples. Same barcodes were used to identify one individual across KDR and mitochondrial libraries so that the three libraries could be ultimately merged by sample. Amplicons tailed with Illumina Nextera® universal sequences were then pooled by column into a single tube and purified using a 0.8-x magnetic beads (SPRIselect, Beckman Coulter) ratio before to perform 15 PCR cycles using Nextera® Index Kit – PCR primers, that adds the P5 and P7 termini that bind to the flow cell and the dual 8 bp index tags. Indexed samples were pooled and quantified by fluorometric quantification (QuantiFluor® dsDNA System, Promega) and visualized on QIAxcel Capillary Electrophoresis System (Qiagen). Libraries were sequenced on a MiSeq run (Illumina) using MiSeq v3 chemistry with 300bp paired-end sequencing.

### Data processing and variant calling

The DDemux program^32^ was used for demultiplexing fastq files according to the P1 barcodes inserted at the 5’-end of each sequence. After demultiplexing, trimmomatic v0.33 was used to discard reads shorter than 32 nucleotides, filter out Illumina adaptor sequences, remove leading and trailing low-quality bases and trim reads when the average quality per base dropped below 15 on a 4-base-wide sliding window. Reads were aligned to two sections of the JAFDOQ010000349 whole genome shotgun sequence with bowtie2 v.2.1.018^33^. The alignment file was converted, sorted, and indexed using Samtools v1.6 and BCFtools v1.8^34^. Coverage and sequencing depth were assessed using bedtools v2.17.0^35^. DNA variants were called using Lofreq 2.1.5^36^ for pooled-mosquito sequencing and Bcftools mpileup callers for single mosquito DNA sequencing, respectively. The bioinformatic pipeline that was used in this work is provided in Supplementary file 2.

### Phylogenetic analyses

Consensus mitochondrial sequences were obtained from aligned bam files using the SAMtools/BCFtools package and seqtk v1.0-r31 (Supplementary file 2). Samples were included in the phylogenetic analysis only if at least 30% of their targeted mitochondrial genome section was covered with a base quality score >20. A background set of 37 full-length mitochondrial genomes were obtained from GenBank^8,9^ to represent the worldwide diversity of *Ae. albopictus* mitogenomes haplogroups. The mitochondrial genome of *Ae. aegypti* (NC_035159) was used as an outgroup in the phylogenetic tree. Consensus sequences were aligned using muscle 5.1^37^ and curated by gblocks software implemented in the seaview version 5.0.4 interface^38^ without option for stringent selection. The curated alignment represented 3,243 nucleotides out of the targeted 4,438 nucleotides (0.73%). It was expanded with eight additional samples harboring a KDR mutations that has between 20% and 30% of their targeted mitochondrial genome section covered with a base quality score >20. The best-scoring maximum-likelihood (ML) tree was generated using this curated alignment with 120 bootstrap replicates with phyml^39^. The GTR nucleotide substitution model was chosen based on the lowest Akaike information criterion (AIC) value using the Smart Model Selection (SMS) in Phyml software^40^. Phylogenetic trees were visualized using the ggtree R package^41^.

### Statistic and data visualization

Descriptive statistics and data visualization were performed in the statistical environment R v4.2.2^42^. Figures were made using the package ggplot2^43^, leaflet^44^, wesanderson color palette^45^, ggtree^41^ and the Tidyverse environment^46^ (Supplementary file 3).

## Acknowledgements

We are deeply grateful to the agents of Altopictus (Flavien Thiers, Renaud Chevalier, Hugo Peyret, *et al.*) and EID Méditerranée (Yves-Marie Kervella and Pascal Eberhart) for their contributions to the field collection and rearing of *Aedes albopictus* samples. We also thanks Igor Filipović for its help with the multiplexing scheme and the creation of the *ddmux* program.

## Author contributions

Albin Fontaine, and Sébastien Briolant designed research. Antoine Mignotte, Guillaume Lacour, Lionel Chanaud and Grégory L’Ambert contributed to the sample collection on the field. Albin Fontaine, Sébastien Briolant, and Nicolas Gomez performed research with the help and supervision of Agnès Nguyen concerning the DNA sequencing. Albin Fontaine and Nicolas Gomez analyzed data. Albin Fontaine and Antoine Mignotte wrote the manuscript with input from all authors.

---

**Supplementary table 1:**
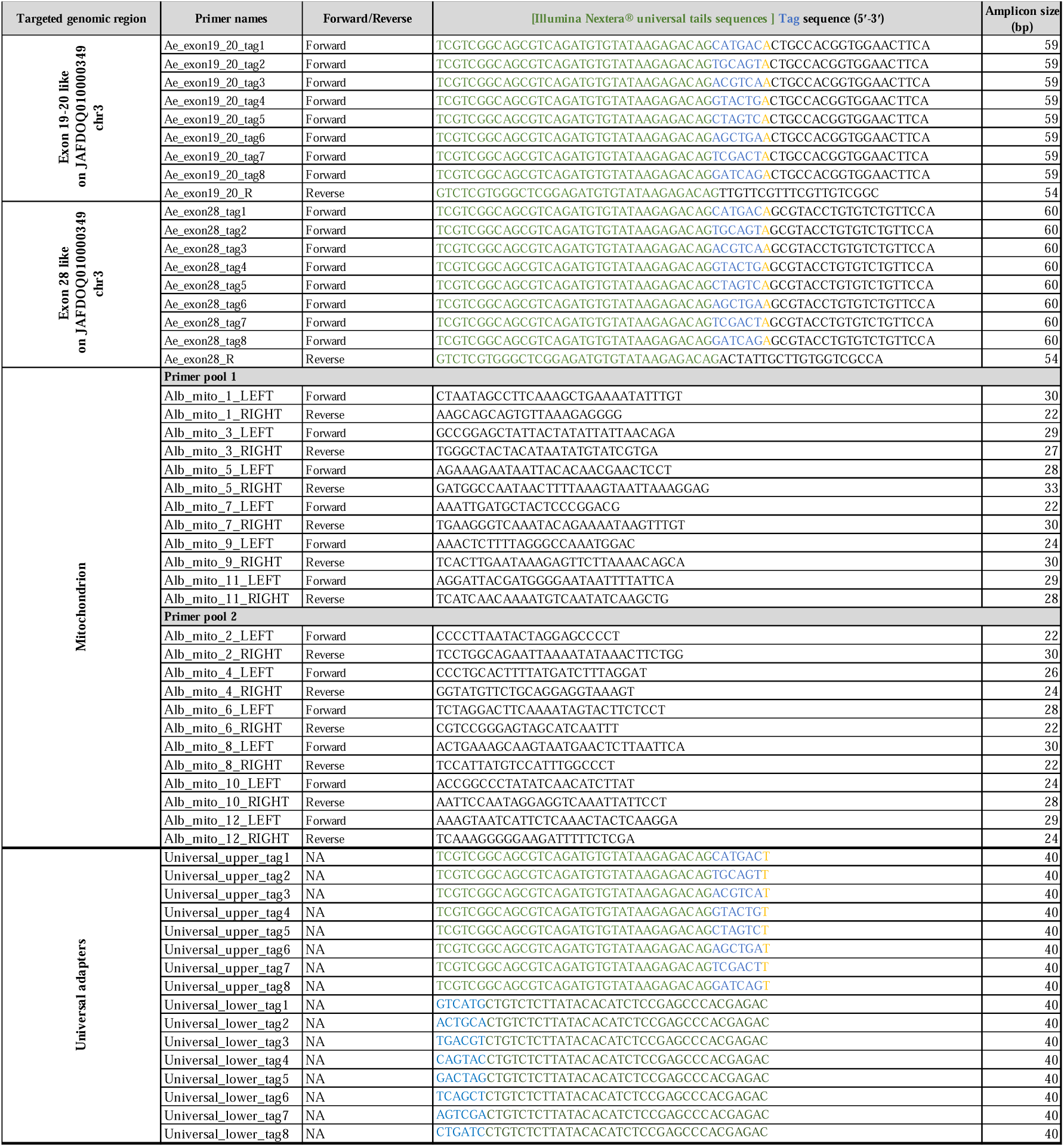
Amplicon-based sequencing systems used in this study to amplify nuclear genomic regions associated with pyrethroid resistance and a 4,438 nucleotides region of the mitochondrial genome of *Ae. albopictus* using ligase-based tiling amplicon sequencing. Primers are presented with their gene targets and amplicon sizes. Illumina Nextera® universal tails sequences are represented in green, and the 6 bp barcodes in blue. An adenine or thymine nucleotides were added to separate barcoded tails from primer sequences. These sequences were directly anchored to primer sequences for KDR amplicons and were added by ligation after amplification for the 12 mitochondrial targets (ligase-based tiling amplicon sequencing).

**Supplementary table 2: List of nonsynonymous mutations revealed by pool DNA amplicon sequencing in *Ae. albopictus* on 4 exons (exon19-like, exon20-like, exon27-like and exon28-like) from the vssc gene.** Mean sequencing quality (QUAL), sequencing depth (DP) and allele frequencies (AF) across samples are indicated for each mutation, with their nucleotide position on our reference and codon position as referred to *Musca domestica* reference genome.

**Supplementary table 3: List of nonsynonymous mutations confirmed by single mosquito DNA amplicon sequencing on 4 exons from the vssc gene.** Sequencing quality (QUAL), sequencing depth (DP) allele frequencies/genotypes (AF, homozygous or heterozygous), and geographic coordinates are represented for each sample with their nucleotide position on our reference and codon position as referred to *Musca domestica* reference genome.

**Supplementary file 1: Interactive map of V1016G/I mutations detected by pool DNA amplicon sequencing.** The map was created with the R leaflet package.

**Supplementary file 2: Bioinformatic pipeline used in data processing and variant calling.**

**Supplementary file 3: R pipeline used in data visualization.**

**Supplementary figure 1:**
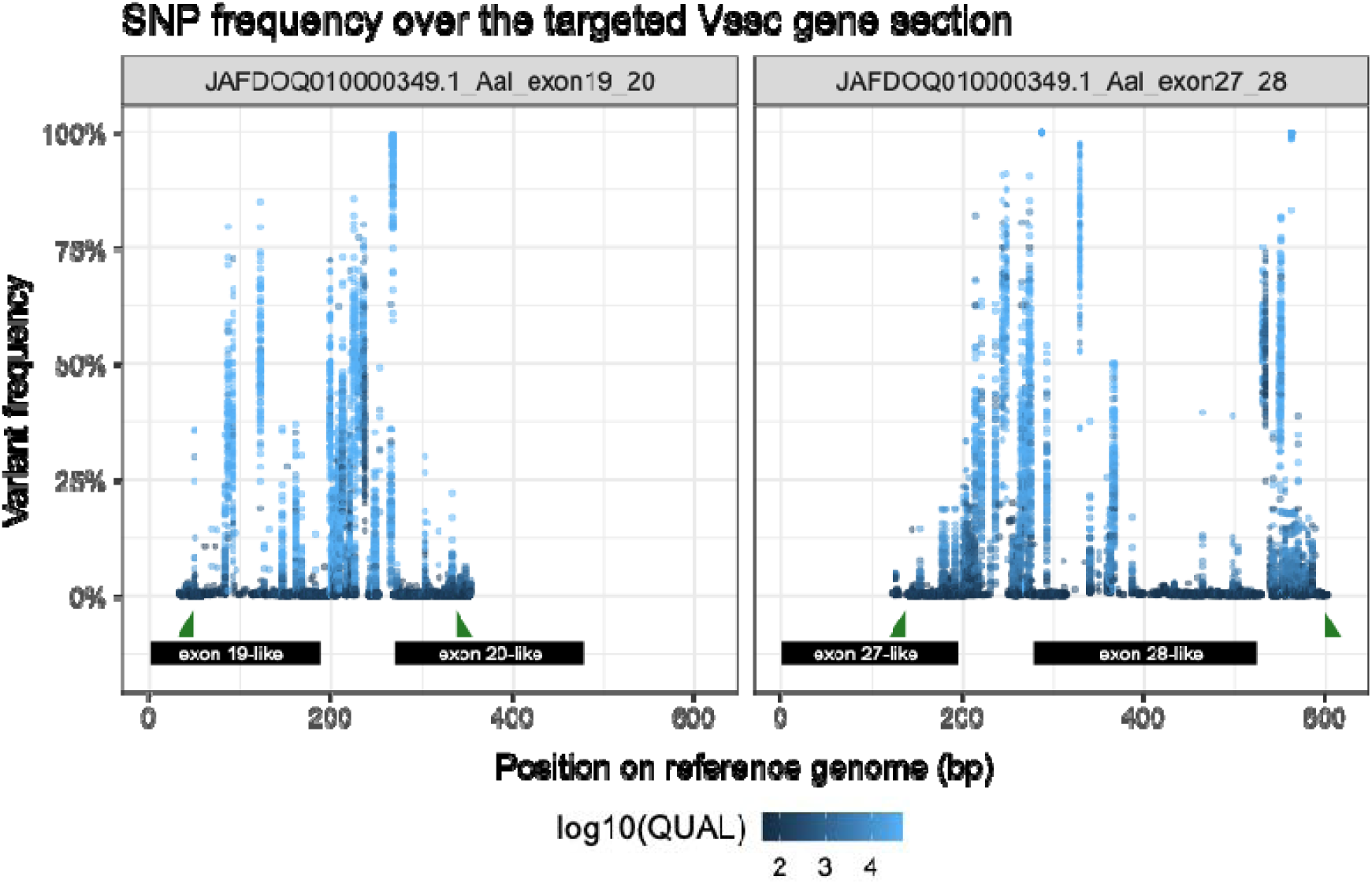
Genetic variant frequencies on two amplified sections of the Vssc gene. Genetic variants are represented with a point colored based on the sequencing quality on a log 10 scale. Exons and primers are represented with black rectangles and green triangles, respectively.

**Supplementary figure 2:**
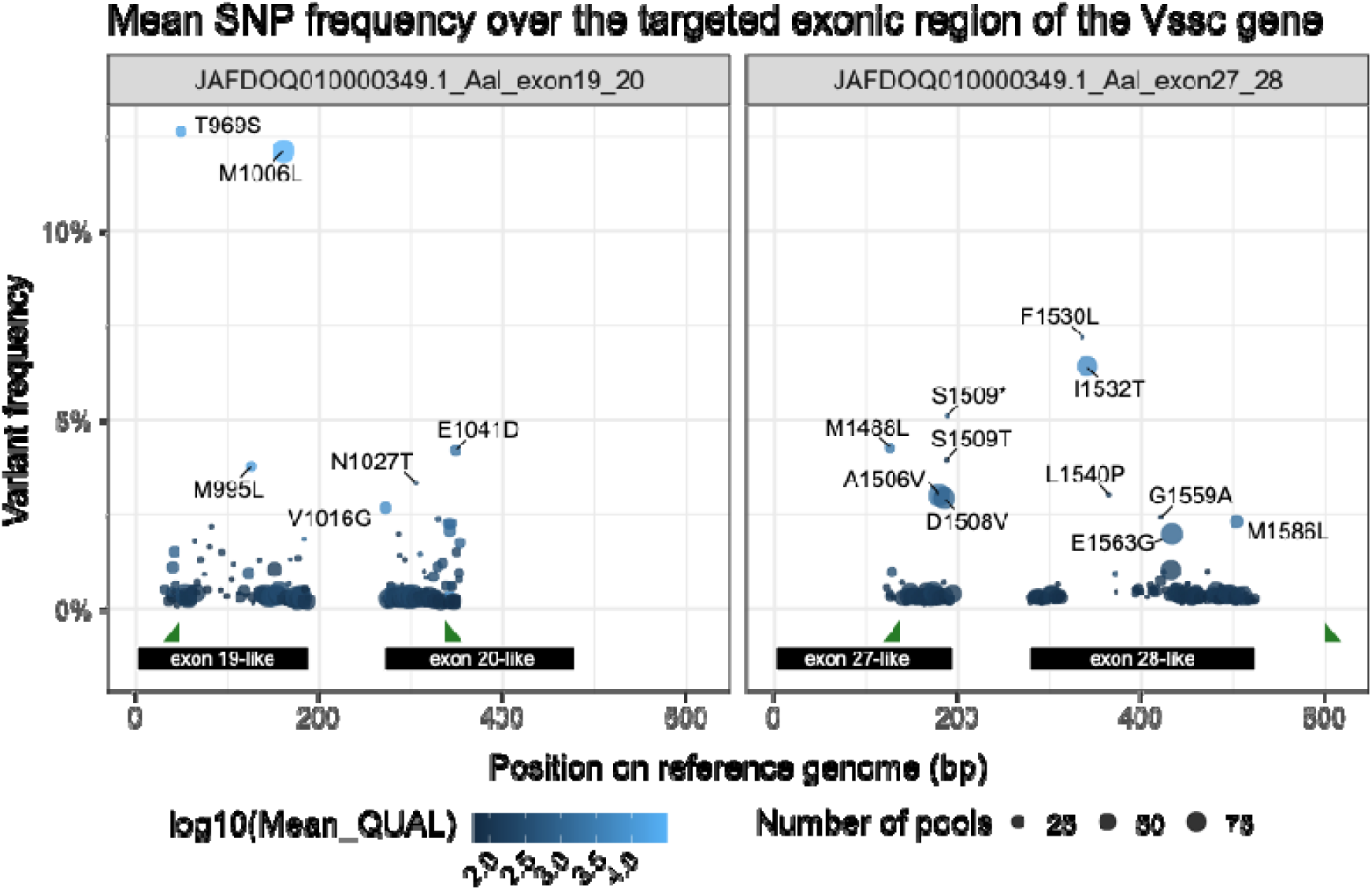
Non-synonymous mutation frequencies on exon of the targeted Vssc gene sections. Genetic variants are represented with a point colored based on the sequencing quality on a log 10 scale and sized according to the number of pools in which the mutation was detected. Mutations are named based on codon positions as referred to Musca domestica reference genome. Only mutations that exceed 2% in frequency are represented to improve the figure clarity. Exons and primers are represented with black rectangles and green triangles, respectively.

**Supplementary figure 3:**
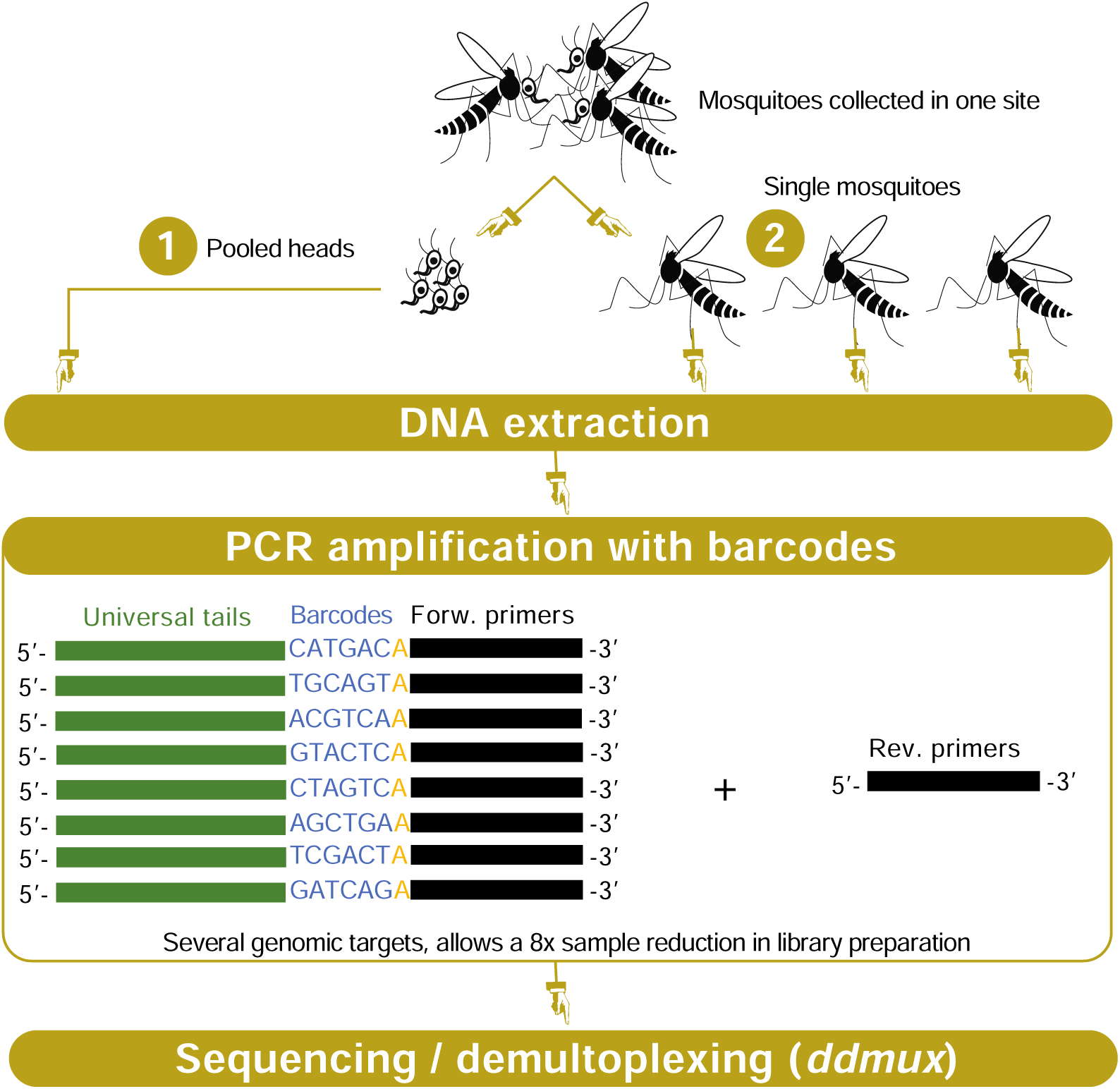
Schematic representation of the multiplexed amplicon-based design. The design allows 8-x sample reduction with 96 samples from one plate being grouped into 12 different tubes, or one plate row, based on eight 6 bp tags integrated in the 5’ end of each amplicon.

